# A combined frequency domain near infrared spectroscopy and diffuse correlation spectroscopy system for comprehensive metabolic monitoring of inspiratory muscles during loading

**DOI:** 10.1101/2023.11.30.569133

**Authors:** Carlos A. Gómez, Laurent Brochard, Ewan C. Goligher, Dmitry Rozenberg, W. Darlene Reid, Darren Roblyer

**Affiliations:** Department of Biomedical Engineering, Boston University, Boston, MA 02125, USA; Keenan Research Centre, Li Ka Shing Knowledge Institute, St. Michael’s Hospital, Unity Health Toronto, Toronto, ON, Canada; Department of Critical Care, St. Michael’s Hospital, Toronto, ON, Canada; Interdepartmental Division of Critical Care Medicine, University of Toronto, Toronto, ON, Canada; Toronto General Hospital Research Institute, University Health Network, Toronto, ON, Canada; Department of Physiology, University of Toronto, Toronto, ON, Canada; Toronto General Hospital Research Institute, Ajmera Transplant Center, University Health Network, Toronto, ON, Canada; Division of Respirology, Temerty Faculty of Medicine, University of Toronto, ON, Canada; Department of Physical Therapy, University of Toronto, Toronto, ON, Canada; KITE – Toronto Rehabilitation Institute, University Health Network, Toronto, ON, Canada

## Abstract

**Significance:** Mechanical ventilation (MV) is a cornerstone technology in the intensive care unit as it assists with the delivery of oxygen in critical ill patients. The process of weaning patients from MV can be long, and arduous and can lead to serious complications for many patients. Despite the known importance of inspiratory muscle function in the success of weaning, current clinical standards do not include direct monitoring of these muscles.

**Aim:** The goal of this project was to develop and validate a combined frequency domain near infrared spectroscopy (FD-NIRS) and diffuse correlation spectroscopy (DCS) system for the noninvasive characterization of inspiratory muscle response to a load.

**Approach:** The system was fabricated by combining a custom digital FD-NIRS and DCS system. It was validated via liquid phantom titrations and a healthy volunteer study. The sternocleidomastoid (SCM), an accessory muscle of inspiration, was monitored during a short loading period in fourteen young healthy volunteer. Volunteers performed two different respiratory exercises, a moderate and high load, which consisted of a one-minute baseline, a one-minute load, and a six-minute recovery period.

**Results:** The system has low crosstalk between absorption, reduced scattering, and flow when tested in a set of liquid titrations. Faster dynamics were observed for changes in blood flow index (BF_i_), and metabolic rate of oxygen (MRO_2_) compared to hemoglobin + myoglobin (Hb+Mb) based parameters after the onset of loads in males. Additionally, larger percent changes in BF_i_, and MRO_2_ were observed compared to Hb+Mb parameters in both males and females. There were also sex differences in baseline values of oxygenated Hb+Mb, total Hb+Mb, and tissue saturation.

**Conclusion:** The dynamic characteristics of Hb+Mb concentration and blood flow were distinct during loading of the SCM, suggesting that the combination of FD-NIRS and DCS may provide a more complete picture of inspiratory muscle dynamics.

## 1. Introduction

Mechanical ventilation (MV) is a lifesaving tool that has become ubiquitous in the intensive care unit (ICU) for critically ill patients with respiratory distress [1]. Pre-COVID-19 pandemic rates of MV in the US were 2.7 episodes per 1000 population and MV use was estimated to cost $27 billion per year [2]. American hospitals reported a 31.5% increase in the number of MV cases during the COVID-19 pandemic [3]. Despite the importance of MV in the ICU, it has several major physiological [4], [5] and psychological [6]–[9] risks, including muscle disuse atrophy, ventilator induced diaphragm and lung injury, post-traumatic stress disorder, depression, anxiety, and cognitive impairments. Thus, it has been recommended that patients should be removed from MV at the earliest opportunity to minimize these risks, especially in older populations [10].

The process of removing patients from MV, also known as the weaning, spans 40% of the duration of MV treatment [11] and starts with a spontaneous breathing trail (SBT) [12]. During a SBT, a patient breathes with little to no assistance of a mechanical ventilator while physicians monitor a wide range of indices. These indices can be categorized as either subjective (i.e. subject displaying signs of pain or difficult breathing) or objective (i.e. heart rate and peripheral oxygen saturation) [13]. Current objective indices help to monitor the state of the patient during SBT but lack crucial insight into the metabolic state of the respiratory muscle themselves. This is unfortunate as the functional capacity of the respiratory muscles is key to the ability for sustain spontaneous breathing. Currently there is no clinical standard to meet this need. While electromyography (EMG) can monitor muscle activation (drive to breathe), it does not indicate the metabolic functional capacity required for spontaneous breathing. Moreover, clinically meaningful analysis of metabolic capacity is not available in real-time and thus is not evaluated as part of clinical practice during weaning. Therefore, there is a critical need for effective non-invasive technologies that can closely monitor patients’ respiratory muscles during weaning in order to guide the readiness and progression of the weaning process and to reduce the duration of MV.

Recently there have been initial investigations, including our own, into the use of near infrared spectroscopy (NIRS) systems to monitor inspiratory muscles during various exercises in healthy subjects, with the stated goal of eventual use for patients on MV [14], [15]. NIRS is a non-invasive optical tool that can measure tissue hemoglobin and myoglobin concentrations via near infrared light. These prior works have investigated the sternocleidomastoid muscle (SCM), a superficial accessory muscle that is recruited during elevated levels of ventilation, including respiratory distress [16]. While NIRS can give insight into the oxygen extraction of tissue, it does not provide a complete metabolic picture as it provides no information about the delivery of oxygen to tissue. When both oxygen extraction and blood flow are measured in unison, these two metrics can be combined via Fick’s principle to calculate the oxygen consumption rate, which may provide a more comprehensive metabolic profile of muscle function [17]. In this work we will describe how we combined a custom frequency-domain NIRS (FD-NIRS) system with a custom diffuse correlation spectroscopy (DCS) system, which can measure blood flow, in order to evaluate metabolic changes of the SCM during a dynamic inspiratory loading protocol. While there have been prior works combining NIRS and DCS systems to measure a range of tissues [17]–[20], this is the first to our knowledge to investigate dynamic changes in inspiratory muscles. It is also the first to characterize the unique alterations in oxygenation, blood flow, and oxygen extraction that occur during inspiratory muscle loading. These results provide an important foundation towards the use of combined NIRS-DCS in the ICU for patients on MV.

## 2. Methods

### 2.1 Custom Combined Diffuse Optical Spectroscopy and Diffuse Correlation Spectroscopy

A custom FD-NIRS system, which has previously been used to monitor the SCM during repetitive quasi-isometric neck flexion in healthy volunteers [15], was integrated with a custom DCS system [21]. Figure 1 shows a block diagram of the combined system and probe layout. The custom fiber probe was fabricated by the Franceshini Group at the Martinos Center at Massachusetts General Hospital. The combined system has three fiber-coupled lasers co-localized via a custom dual source and single detector fiber probe. The FD-NIRS system has both a 730 nm and an 830 nm laser (Blue Sky FMXL730-030YFGA and Thorlabs LPS-830-FC), which are modulated by the direct digital synthesizers (DDS) at 139 and 149 MHz frequencies respectively. The DCS system uses a long coherence laser (CrystaLaser DL852-100-S) with a wavelength of 852 nm. The DCS laser is coupled to 105 µm core fiber (Thorlabs FG105LGA) with a numerical aperture (NA) of 0.22 that is split between two prisms, each 3.5 mm x 3.5 mm, with one prisms receiving 75% of the illumination power and the other receiving the remaining 25% of the illumination power. Each FD-NIRS laser is coupled to a separate 400 µm core fiber (Thorlabs FT400EMT) with NA 0.39. These fibers are coupled to the aforementioned prism receiving 25% of the DCS illumination power. The use of dual prisms allows the illumination power to be distributed across a larger skin area, thus enabling overall higher illumination optical power while staying within American standard safety institute limits.

**Figure 1.**
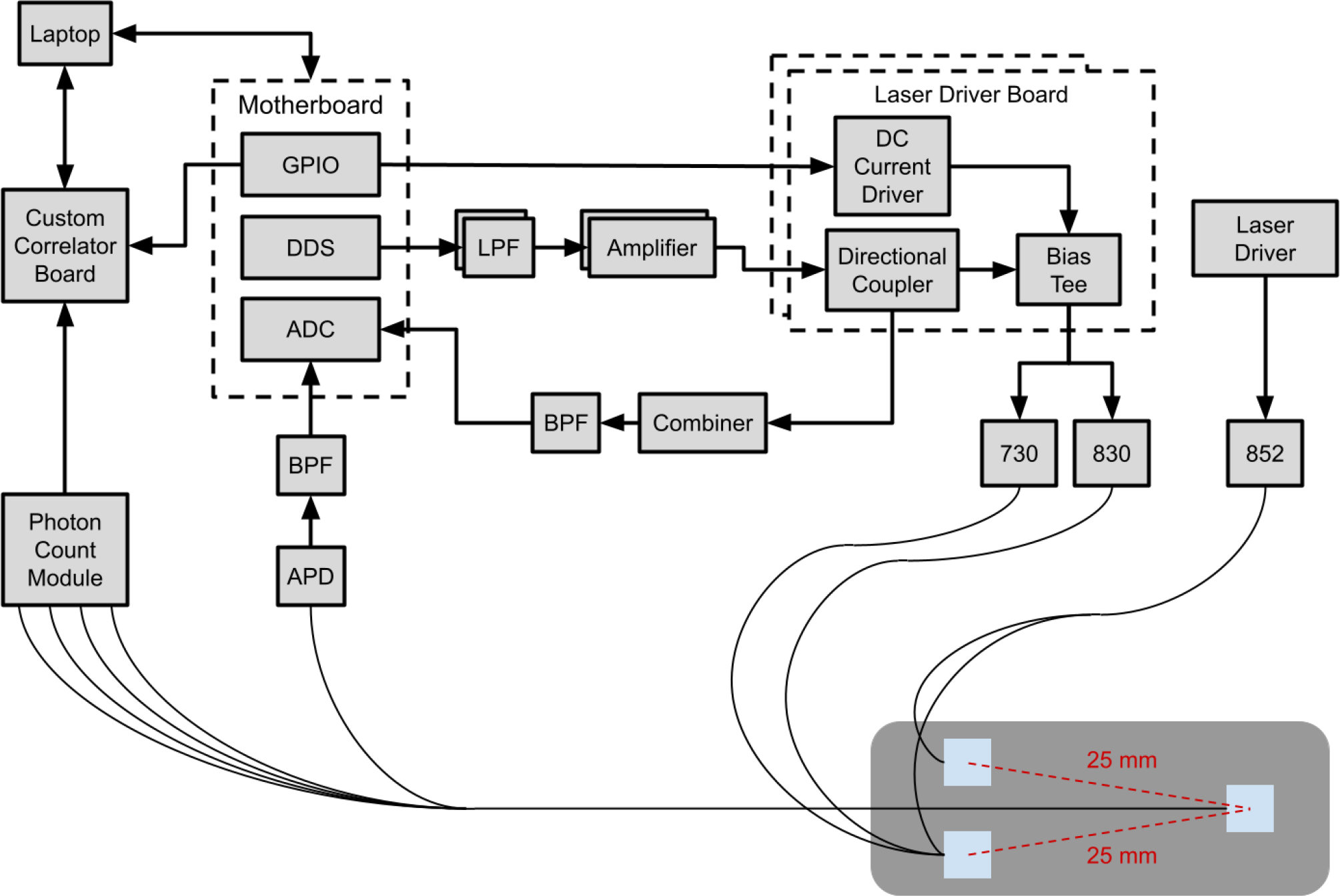
Block diagram of the custom combined FD-NIRS and DCS system. Both systems are controlled by a single computer via separate custom software. The FD-NIRS system is composed of a custom motherboard that contains three daughter boards: an analog to digital converter (ADC) board, a direct digital synthesize (DDS) board, and a general purpose input and output (GPIO) board. The DDS board outputs two sinusoidal signals that first travel through a lowpass filter (LPF) with a frequency cutoff at 400 MHz, which then is amplified before being combined with the direct current (DC) via a bias tee. A reference signal for each FD-NIRS laser is transmitted to the ADC board by having a directional coupler send a signal to a combiner that combines both signals into one and goes through a bandpass filter (BPF) with a frequency range of 110 – 180 MHz. The modulated signals with DC offsets drives the two lasers at two separate frequencies (139 and 149 MHz) which are optically coupled to the DCS’s long coherence laser by a custom probe. Once the FD-NIRS optical signal has traveled through the tissue the signal is detected by an avalanche photon detector (APD) and the signal goes through the same BPF as in the reference signal pathway before going to the ADC board. Additionally the signal from the long coherence laser is detected by a photon module that contains four separate detectors and the signal is sent to a custom correlator board that time stamps the signal. The systems are temporal multiplexed by having the FD-NIRS lasers modulated on and off which is done via the GPIO board of the FD-NIRS. Additionally, the GPIO board sends a signal to the correlator board in order to time stamp when the FD-NIRS lasers were on. A custom probe with two source 3.5 mm prisms (blue squares) was used to split the DCS light source in order to increase the signal. Additionally, the probe only had a single detector 3.5mm prism that co-localized both NIRS and DCS signal.

An avalanche photodiode (APD) (Hammatsu S11519-30) is used as the detector for the FD-NIRS system and is coupled to a fiber bundle with NA 0.66. A single photon-counting module (Excelitas Technologies SPCM-AQ4C) is used for the DCS system and is fiber coupled to 125 µm core fiber (Thorlabs 780HP) with NA 0.13. Both fibers are coupled to the same 3.5 mm x 3.5 mm detector prism. The source detector separation of the custom probe is 25 mm. The systems were temporally multiplexed and had a sampling rate of 2 Hz, but down sampling was performed in post processing to increase the signal to noise ratio of the DCS system, resulting in an effective sampling rate of 0.5 Hz for the combined system. Custom software was used to control both the FD-NIRS and DCS system via the same laptop.

Tissue concentrations of oxygenated hemoglobin plus myoglobin (oxy [Hb+Mb]) and deoxygenated hemoglobin plus myoglobin (deoxy [Hb+Mb]) were determined using measurements from the FD-NIRS system. This was done by comparing the reference signal from the DDS and APD signal. Changes in the amplitude and phase induced by the tissue were calculated by a field programmable gate array in the digital FD-NIRS electronics. The information was then fed into a single layer look up table (LUT) in order to recover both absorption and reduced scattering coefficient (µ_a_ and µ’_s_) of the tissue [22]. The LUT was generated using Monte Carlo (MC) simulations that assumed an index of refraction of 1.37 [23] and an anisotropy value of 0.9 [24]. A calibration procedure was performed in order to remove the instrument response function [25]. The recovered µ_a_ from both wavelengths was then fed into the Beer’s Law using known chromophore extinction coefficients to recover both Oxy [Hb+Mb] and Deoxy [Hb+Mb]. An assumption of 20% lipid fraction and 62.5% water fraction was used for the tissue [26]–[29]. Total [Hb+Mb] was calculated by adding Oxy [Hb+Mb] and Deoxy [Hb+Mb] together. Tissue saturation (S_t_O_2_) was then derived from oxy and deoxy [Hb+Mb] via the following equation:

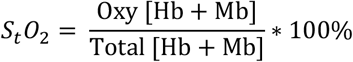

Blood flow index (BF_i_) was determined using measurements from the DCS system. The custom correlator board time stamped the photon signal from the single photon counting module and the signal was autocorrelated with itself over a small time period; this term is known as the intensity autocorrelation curve (g2). The g2 and the µ_a_ and µ’_s_ from the FD-NIRS system were fed into a single layer LUT [30], generated using MC simulations that had an assumed index of refraction of 1.37 [23] and an anisotropy value of 0.9 [24], in order to recover the BF_i_. The oxygen metabolic rate of (MRO_2_) was then derived by the following equation based on Fick’s principle:

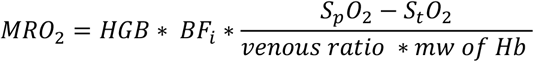

Where HGB is the hemoglobin concentration of blood, SpO_2_ is the peripheral arterial oxygen saturation, the *venous ratio* is the proportion of blood volume in the venous circulation, and *mw of Hb* is the molecular weight of hemoglobin. The following assumptions were made for these parameters based on prior literature. HGB values of 14 g/dL and 16 g/dL were assumed for females and males respectively [31], an SpO_2_ of 98% was assumed [32], a *venous ratio* of 0.75 was assumed [33], and the *mw of Hb* of 64,500 g/mol was assumed [31].

### 2.4 Cross Talk Evaluation

Three different liquid phantom titrations were performed to assess the cross talk between the three core measured parameters (µ_a_, µ’_s_, and BF_i_). All titrations were performed in a container with the following dimensions: 150 mm x 95 mm with 70 mm depth. The optical probe was placed directly on the surface of the liquid. For the absorption titration, an initial intralipid solution of 0.5% lipid was created by diluting 20% stock intralipid with deionized water. For each titration step, 0.48 mL of the batch solution was removed from and replaced with 0.48 mL of nigrosin solution with 1.5 g/L concentration that was composed of nigrosin diluted in 0.5% intralipid. Between each titration step, the solution was mixed for 60 seconds with an additional 90 second pause before measuring in order to ensure there was only Brownian motion in the solution. Similarly, for the scattering titration, an intralipid solution of 0.39% was created and for each titration step 2 mL was removed from the batch solution and 2 mL of 20% intralipid added. Again, the solution was mixed for 60 seconds with a 90 second pause before measuring. For the flow titration, a 0.35% intralipid solution was used and it was constantly stirred with a magnetic stirrer at 64 rpm. Each titration step involved increasing the speed of the stirrer by 7 rpm and waiting 60 seconds before measuring. Cross talk was defined as the ratio of the normalized to baseline undesired change to the desired change expressed in decibels.

### 2.5 Healthy Volunteer Study

All measurements were conducted under an institutionally approved protocol (BU IRB 5618E). FD-NIRS and DCS measurements were conducted on 14 healthy volunteers (7 females and 7 males) aged 26.3 ± 1.4 years while they performed a breathing exercise with a respiratory device(s) (Philips Threshold IMT, POWERbreathe Plus IMT – Light Resistance, and POWERbreathe Plus IMT – Medium Resistance). First, subjects performed three maximum inspiratory pressure (MIP) tests and their values were recorded from a pressure gauge (Vacumed 1505-120 Respiratory Pressure). The mean MIP was used to determine a high load (90% of MIP) and a moderate load (30% of MIP) for each subject. While the subjects were sitting upright, their right side SCM was located by having them look down and then to the left, which caused the SCM to be visible to an operator. The custom probe was placed over the SCM at approximately the center of the muscle while the subject was at a neutral head position; the specific location over the SCM was chosen to maximize the signal from the two instruments. Subjects preformed two eight-minute breathing exercises that consisted of one minute for baseline, one minute for load, and six minutes of recovery. During the eight minutes, subjects breathed only through their mouth during baseline and recovery, and breathed through the respiratory device during the load phase. Subjects were given a five second count down before both the start and end of the load phase. Each subject performed a moderate load measurement first and subjects were given a 10-minute break before the start of the high load measurement.

### 2.6 Data Processing

Time traced of six extracted parameters (BF_i_, Oxy [Hb+Mb], Deoxy [Hb+Mb], Total [Hb+Mb], S_t_O_2_, and MRO_2_) were filtered through a second order Butterworth low pass filter with frequency cutoff 0.02 Hz to remove breathing oscillations. Offset time and percent change metrics were then extracted for both loads of each subject from the filtered time traces (Figure 2). There was a wide range of responses to the load, with the most common being a double hump trace, thus two regions of activation were denoted. Offset was always calculated from the start of the load (t = 60 seconds) due to the fact some subjects had only one peak which appeared in either region of activation. While Figure 2 shows peaks during activation, for some parameters (i.e., deoxy [Hb+Mb]) there was a decrease response so the valleys were selected in those time traces. Baseline values were calculated by averaging the initial 50 seconds of the filtered time trace in order to avoid any anticipatory response. Normalization was performed by dividing the extracted parameter time traces by the baseline value. Respiration rates for baseline and both regions of activation were calculated by first filtering the time trace from the amplitude FD-NIRS signal at 730 nm with a second order Butterworth high pass filter with frequency cutoff 0.03 Hz. The filtered data was then used to find the mean time difference between peaks of each breath in a 30 second time window, this period between breaths was used to calculate breaths per minute by dividing 60 second by the period. Statistical analysis was done by running unpaired two-tailed Student’s t-test to compare various extracted metrics (offset, normalized to baseline, the first 50 seconds, percent change, and respiration) between sex, load, and regions of activation. Systematical testing was done in order to determine if sex, load, and regions of activation had any statistically significant effects on the extracted metrics.

**Figure 2.**
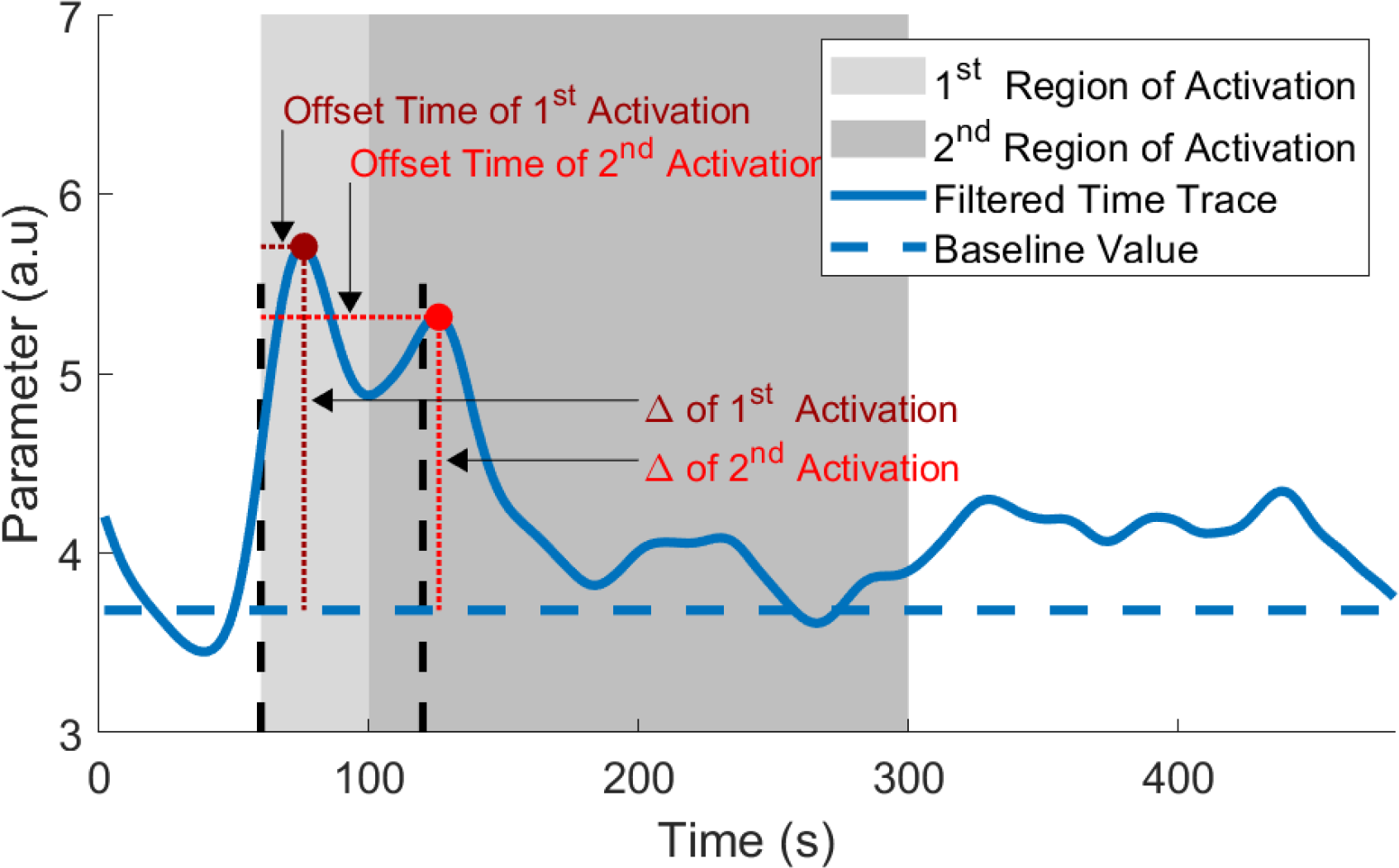
Example filtered time trace with offset and percent change from baseline (Δ) for the two regions of activation during the eight-minute breathing exercise. Vertical black dashed lines indicate the start and end of the one-minute load portion of the respiratory exercise. Baseline value was determined by averaging the initial 50 seconds to avoid anticipatory effects. Offset time was determined by calculating the time difference from the event to the start of the load at the 60 seconds, which is indicated by the first vertical black dashed line. Percent change from baseline was calculated by measuring the difference from the event’s value to baseline.

## 3. Results

### 3.1 Cross Talk

The titrations results are shown in Figure 3. Each had a low (< -11 dB) crosstalk between the three measured parameters. For the absorption titration, absorption increased by 544% and 286% for 730 nm and 830 nm respectively while reduced scattering increased by only 13% and -3% for 730 nm and 830 nm respectively and BF_i_ increased by 19%. For the scattering titration, reduced scattering increased by 70% and 67% for 730 nm and 830 nm respectively while absorption decreased by only 3% and 2% for 730 nm and 830 nm respectively and BF_i_ increased by 1%. For the flow titration, BF_i_ increased by 276% while absorption changed by only -8% and 1% for 730 nm and 830 nm respectively and reduced scattering increased by 0% and 0% for 730 nm and 830 nm respectively. Absorption and scattering had small variance in all three titrations, as seen by the error bars in Figure 3. On the contrary, BF_i_ had larger variance, especially in the flow titration, which most likely arose from the increase in rpm of the stirrer. The increase in speed reduced the stability of stirrer as larger fluctuations in rpm speed were noted at higher speeds, which likely explains the increase in error bar size at higher speeds in Figure 3.

**Figure 3.**
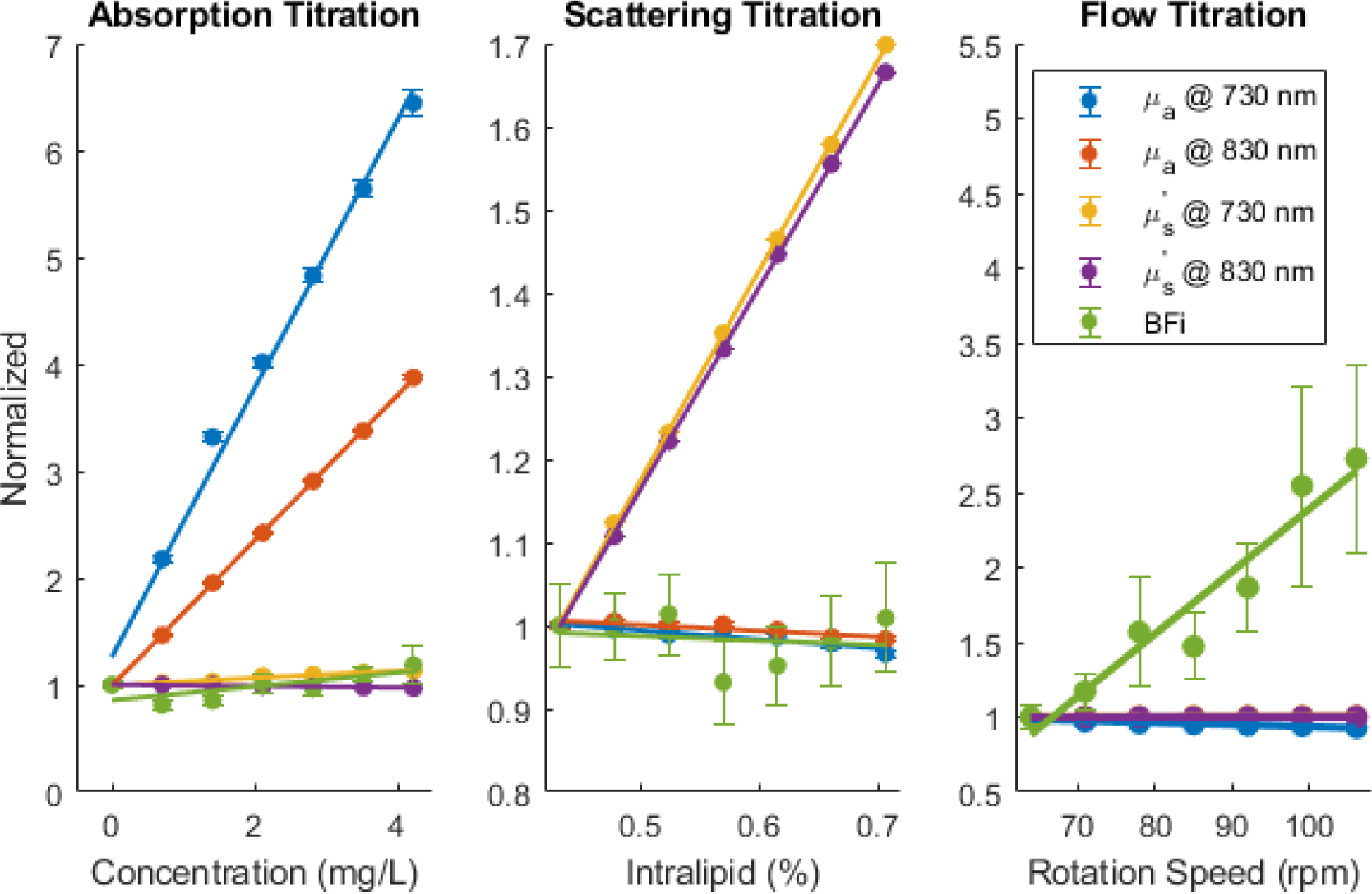
Normalized change from baseline of µ_a_ and µ’_s_ from the FD-NIRS system and BF_i_ from the DCS system during each of the three titrations. Error bars at each titration step represent the standard deviation. Solid line represents best linear fit curve. Across all three titrations, there was low cross talk between measure parameters. The absorption titration (left graph) had on average a 415% increase in µ_a_ while only having on average a 5% increase in µ’_s_ and 19% increase in BF_i_. The scattering titration (center graph) had on average a 68.5% increase in µ’_s_ while only having on average a 2.5% in µ_a_ and a 1% increases in BF_i_. The flow titration (right graph) had a 276% increase in BF_i_ while on average a 3.5% decrease in µ_a_ and no change in µ’_s_. All titrated parameters had a linear response during its titration as seen by the best fit trends. Both µ_a_ and µ’_s_ had minor variation in values as seen by the small error bars while BF_i_ had larger variation.

### 3.2 Healthy Volunteer Study

Baseline values were calculated for all six parameters for female and male participants and the results can be seen in Supplementary Table 1. Only oxy [Hb+Mb], total [Hb+Mb], and tissue saturation had significant difference in baseline values between the sexes with p values of 0.011, 0.018, and 0.044 respectively.

Figure 4 shows the mean of all males (n=7) time traces for Oxy [Hb+Mb], BF_i_, and MRO_2_, and helps to highlight the most common features observed in the data. For example, the time offset values were typically shorter for BF_i_ and MRO_2_ compared to hemoglobin concentration and saturation changes. Additionally, the percent changes were typically larger for BF_i_ and MRO_2_ compared to hemoglobin changes. A double hump feature was commonly observed after the start of the load as shown by the two peaks in the measured parameters.

**Figure 4.**
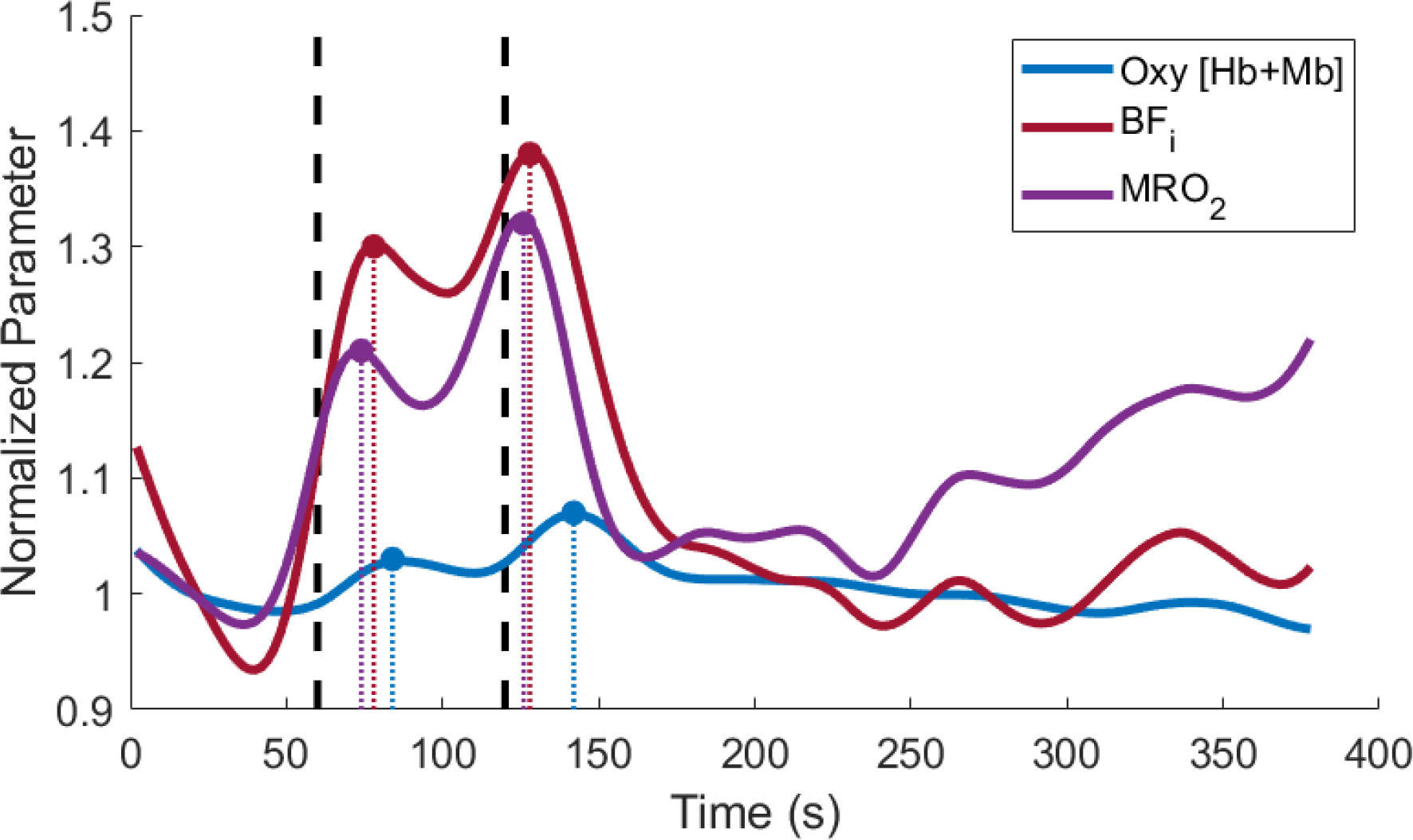
Time traces of the mean of all male data (n=7). Traces were normalized to baseline (i.e. the first 50 seconds). Vertical black dashed lines indicate the start and end of the one-minute load portion of the respiratory exercise. Data has the last 100 seconds removed due to one subject coughing which resulted in large muscle activation. The time offset values (i.e., time from the start of the exercise, indicated by the first vertical dashed line), were typically shorter for BF_i_ and MRO_2_ compared to hemoglobin + myoglobin concentration and saturation changes. Additionally, the percent changes were typically larger for BF_i_ and MRO_2_ compared to hemoglobin + myoglobin changes.

The mean filtered time traces for all subjects for the six parameters are plotted in Figure 5, and all individual filtered time traces can be seen in supplementary Figures 1 – 6. The time traces were separated by sex and by load resulting in four mean time traces per subplot. Activation of the SCM was observed in all six metrics as indicated by changes from baseline after the start of the load, with deoxy [Hb+Mb] being the only metric to show a decrease from baseline while the other five metrics had increases from baseline. Additionally, the mean time traces showed a clear double hump feature within the 240-second time window after the start of the load. These double features led us to extract offset and percent change for the two regions of activation for all six parameters. The mean and standard deviation for the offset for all sex and load combinations during both regions of activation can be seen in Supplementary Table 1. Additionally, the mean and standard deviation for the absolute change for all sex and load combinations during both regions of activation can be seen in Supplementary Table 2. The rate of respiration for all sex and load combination during baseline and both regions activation can be seen in Supplementary Table S3. There was no significant difference for the respiration rate between sex, load, and region vs baseline.

**Figure 5.**
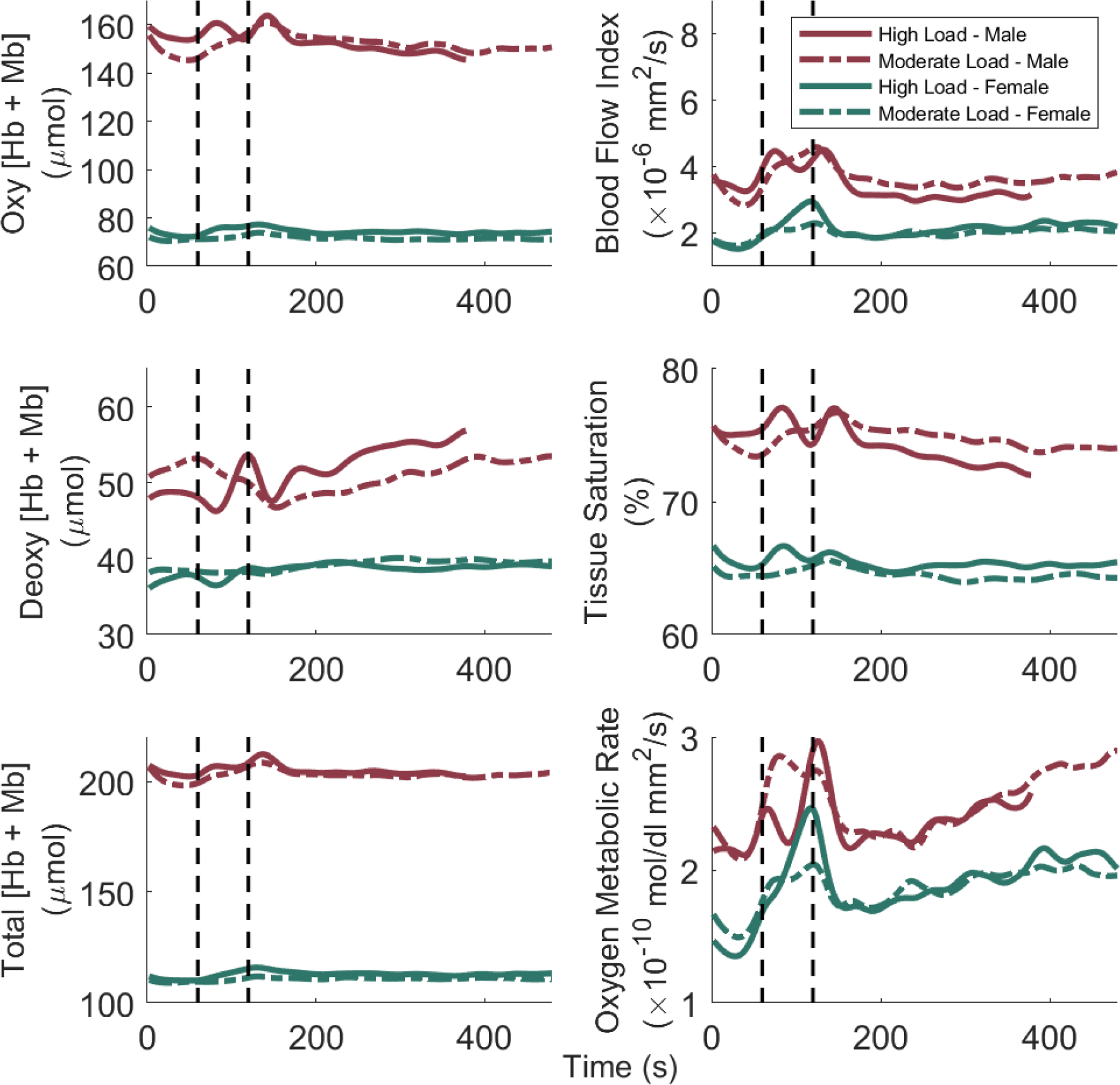
Mean filtered time traces of the six muscle parameters from the combined system. Vertical black dashed lines indicate the start and end of the one-minute load portion of the respiratory exercise. The high load, male time traces have the last 100 seconds removed due to one subject coughing which resulted in large muscle activation. Only the oxy and total [Hb+Mb] had statistical differences between sexes due to large subject variances. The majority of mean time traces had a double perturbation event with most having a positive change from baseline with only deoxy [Hb+Mb] having a negative change from baseline. The offset time of the second perturbation for MRO_2_ was significantly different between the sexes (p = 0.03).

The offset values from the six muscle parameters were analyzed for each sex and region of activation to determine if there were any differences between the temporal dynamics of the six parameters (Figure 6). The offset values were separated by sex and not by load due to the fact that there was no significant difference between the loads. There was, however, a significant difference between the male and female offsets (p = 0.03) for the second region of activation of BF_i_ during high loads. Females showed no significant differences between the offsets of the six parameters for either region of activation. Males had significantly shorter offsets for both BF_i_ (p = 0.043, p = 0.045, and p = 0.0008) and MRO_2_ (p = 0.046, p = 0.046, and p = 0.009) in the second region of activation when compared to oxy [Hb+Mb], deoxy [Hb+Mb], and total [Hb+Mb].

**Figure 6.**
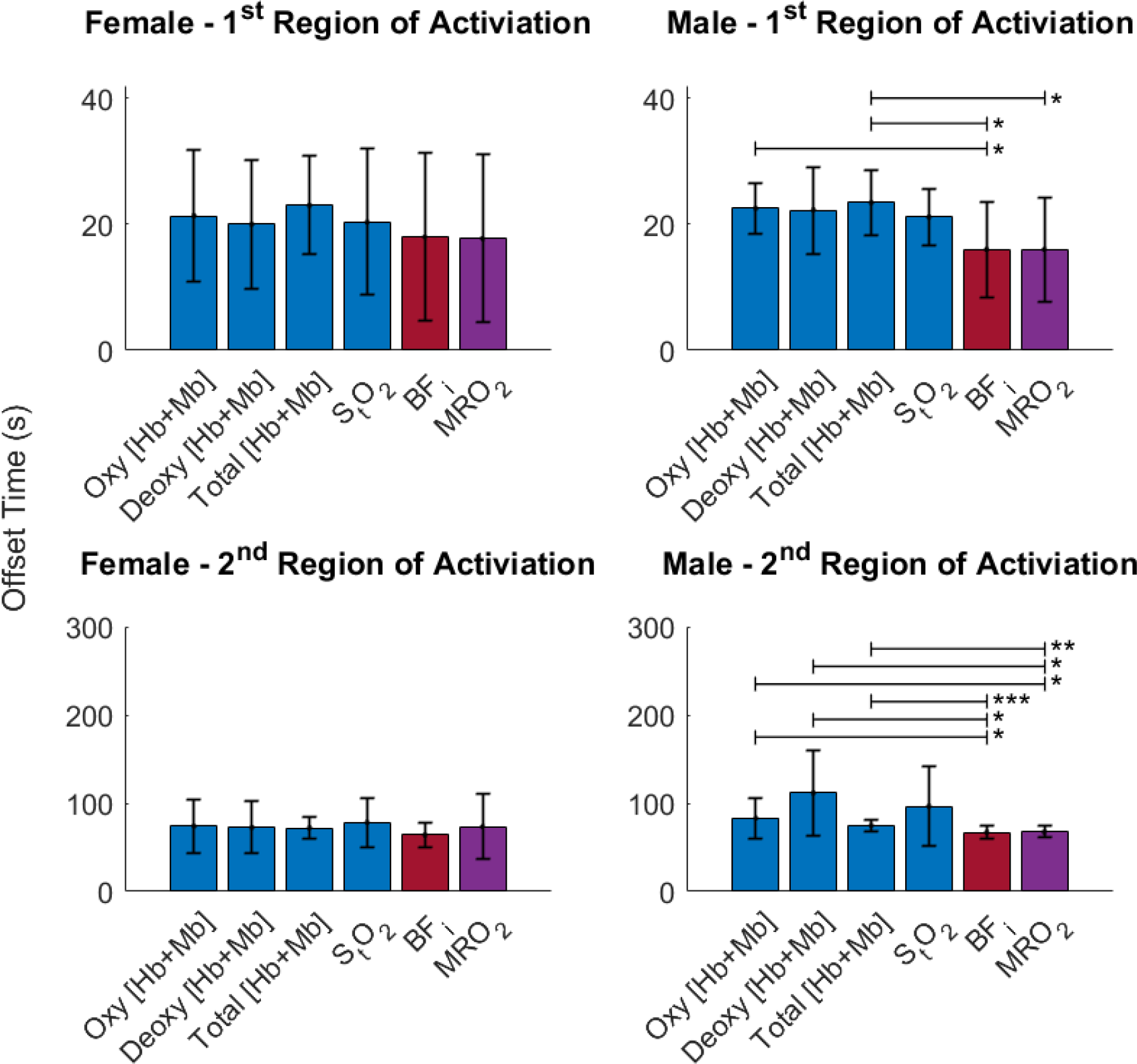
Offset values for the six muscle parameters. Both moderate and high loads are included. Males had significantly shorter offsets for both BF_i_ and MRO_2_ when compared to total [Hb+Mb], oxy [Hb+Mb], and deoxy [Hb+Mb]. There was no statistically significant differences for females. For males, these results suggest that MRO_2_ is tightly coupled with the BF_i_ temporal responses. * p < 0.05, ** p < 0.01, *** p < 0.001,

Similar statistical analysis was performed on percent change of the six parameters (Figure 7). Data was pooled as there was no significant difference between sex, load, or region of activation. BF_i_ and MRO_2_ had the largest increase compared to the other metrics. Additionally, oxy [Hb+Mb], total [Hb+Mb], and S_t_O_2_ had increases from baseline while a decrease from baseline occurred in deoxy [Hb+Mb].

**Figure 7.**
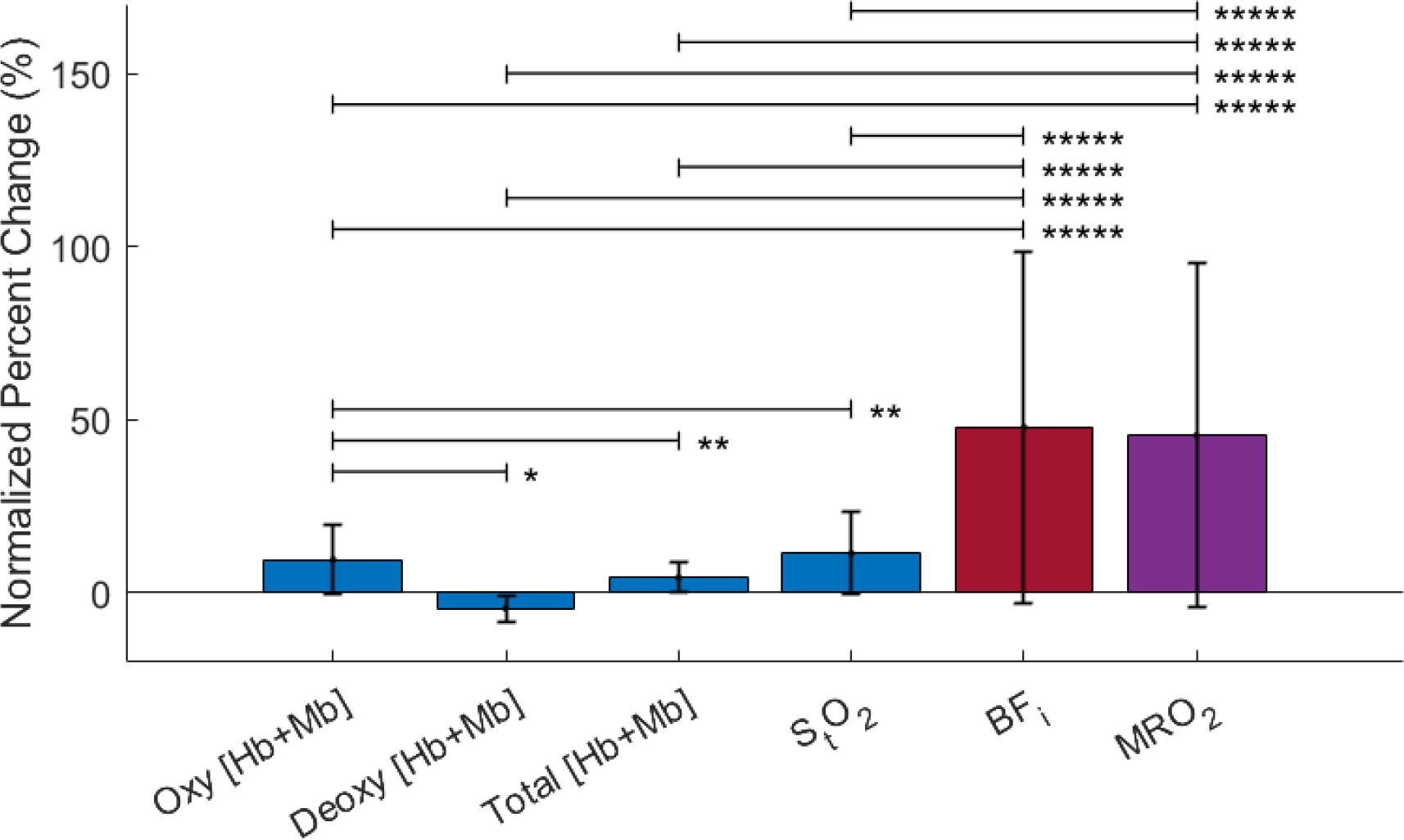
The percent change for the six muscle parameters when pooled across sex, region of activation, and load. Data was normalized to baseline (i.e. the first 50 seconds). BF_i_ and MRO_2_ had the largest percent increases. Deoxy [Hb+Mb] decreased while oxy [Hb+Mb] increased and total [Hb+Mb] only increased slightly. An increase in tissue saturation was also observed. * p < 0.05, ** p < 0.01, *** p < 0.001, **** p < 0.0001, ***** p < 0.00001

## 4. Discussion

We have successfully combined a custom FD-NIRS system and custom DCS system to operate in unison for the purpose of dynamically monitoring the SCM during loading. The combined system was validated via three liquid titrations and a healthy volunteer study. The titrations showed minimal crosstalk in the measured parameters (µ_a_, µ’_s_, and BF_i_) when swept across anticipated physiological ranges. Additionally, the changes in measured parameters (oxy [Hb+Mb], deoxy [Hb+Mb], total [Hb+Mb], StO2, BF_i_, and MRO_2_) from the SCM were continuously monitored during respiratory loading via a respiratory device. These parameters were analyzed to determine the typical physiological response of the SCM during a short perturbation load in young healthy subjects. Several key trends were observed, including shorter time offsets for BF_i_ and MRO_2_ compared to hemoglobin + myoglobin based parameters in males, and larger percent changes in BF_i_, and MRO_2_ compared to hemoglobin + myoglobin based parameters in both males and females. These data suggest that, at least in some circumstances, the dynamic characteristics of blood flow and oxygen extraction are substantially different during SCM loading compared to NIRS-based hemoglobin + myoglobin measures. This suggests that the use of combined FD-NIRS and DCS may provide a more complete picture of inspiratory muscle dynamics, potentially assisting in the evaluation of patients under MV in the future.

The key finding that BF_i_ and MRO_2_ had larger changes than hemoglobin + myoglobin based parameters was also observed in a prior study during the initial response of a quadriceps exercise [17]. This, combined with the fact that BF_i_ and MRO_2_ had faster offset times compared to hemoglobin + myoglobin based parameters in males, is consistent with the hypothesis that a change in oxygen delivery (through increased blood flow via vasodilation mechanisms), rather than a change in oxygen extraction, is the more rapid and dominant metabolic response mechanism for the type of inspiratory perturbation evaluated here. It is of note that larger changes in oxy [Hb+Mb] and deoxy [Hb+Mb] have been reported when using standalone NIRS system when measuring the SCM [14], [15]. This difference may be due to the different type of muscle activation utilized that recruited more motor units due to higher loads [14], [15] and longer duration [14] in this study compared to the prior work. This study used a short one-minute load, whereas the prior studies loaded the SCM until failure. Short activation, like the one-minute load used here, might be more feasible as a means to evaluate readiness for weaning in an ICU setting as it is more rapid and may be less likely to cause respiratory muscle fatigue or damage. Additionally the short load period in this study contributed to there being no statistically significance between the moderate and high loads. There was no evidence to show that the subjects had reach their max oxygen consumption rate as the MRO_2_ did not show signs of plateauing during either region of activation which could have lead similar temporal and percent change values between loads.

This study showed some differences between the sexes. This is consistent with previous work that has shown differences in respiratory system mechanics during activation between the sexes [34]. Baseline values in oxy [Hb+Mb] and total [Hb+Mb] were different between the sexes, with males having higher concentrations on average, which has also been reported by other groups [15], [19], [35]. This may be attributable to the known hemoglobin content difference between sexes [36]. There were also some sex dependent responses in temporal offset of some parameters. For example, during high loads in the second region of activation, statistical differences were observed in offset in BF_i_ between the sexes. Overall males had more rapid BF_i_ and MRO_2_ activation in both regions of activation while females did not have any single parameter change faster than the rest. These offset trends between the sexes are somewhat difficult to interpret as the results could have been impacted by several factors: short load period, anticipatory response, and subject variance in respiratory muscle group response.

The mean time traces showed a unique double hump in all six-muscle metrics, although there was significant variance between subjects. This variability most likely arises from the fact that SCM is only a single muscle in a group of muscles that work together to allow inspiration to occur. When faced with a load the body might respond by activating other muscles of inspiration including the diaphragm, intercostals, and scalene muscles, in various proportions alongside the SCM. Further work should investigate the subject variance response by probing various respiratory muscles during similar loads. Additional work should investigate the respiratory response of the SCM during extended respiratory loading.

We showed here that the SCM oxygen consumption is driven by an increase in blood flow for a short period of load, but this might not be the case for extended loads. Work with FD-NIRS and DCS on the quadriceps during exercise have shown that blood flow dominates the early stages of muscle activity but drops in tissue saturation do occur near the muscle failure point [17]. The SCM most likely responds in a similar manner and this could also be insightful to compare between healthy subjects and patients on MV with or without established respiratory muscle weakness.

Going forward the combined methodology of FD-NIRS and DCS may be useful in characterizing inspiratory muscle response during weaning from MV, with the eventual goal of noninvasive identification of patients who are ready for weaning. Additionally, the technique might also play a key role in timing of intubation for MV treatment. Further study in healthy volunteers and patients on MV is warranted.

## 5. Conclusion

Custom FD-NIRS and DCS devices were combined to continuously monitor the SCM during a one-minute task of respiratory exercise. There were minor differences between the sexes in some baseline parameters. Importantly, the proportional increases in BFi and MRO_2_ were greater than changes in hemoglobin + myoglobin based parameters for all subjects, and the temporal dynamics of BFi and MRO_2_ were faster in males compared to hemoglobin + myoglobin based parameters. These trends suggest that metrics measured with FD-NIRS and DCS have distinct dynamics during loading in the SCM, and therefore it may be beneficial to utilize both technologies in the future when monitoring patients on MV.

## Disclosures

The authors report no conflicts of interest.

## Code, Data, and Materials Availability

Scripts, data, and associated instructions for performing each step of the analysis from raw data up till filtered time traces of each subject are provided in a repository on Github: https://github.com/BU-BOTLab/FDNIRS_paper_JBO2023. Furthermore, the exact time points and perturbation values from all parameter for each subject are included in the respiratory. Statistical analysis are performed on their recorded values and further information can be provided upon requested.

## Acknowledgements

We would like to thank Lina Lin for helping in printed circuit board design and troubleshooting, Nikola Otic for help in building and training on the DCS system, Bernhard Zimmerman for help in building the DCS system, Zachary Starkweather for help in building a custom probe, and Mari Franchesceni and David Boas for their insight on DCS.

## Funding

The authors gratefully acknowledge funding from the NIH NIBIB award R21EB031250 and the NIH QBP fellowship T32GM145455. Additionally, this material is based upon work supported by the National Science Foundation Graduate Research Fellowship Program under Grant No. 2234657. Dmitry Rozenberg receives research salary support from the Sandra Faire and Ivan Fecan Professorship in Rehabilitation and Temerty Faculty of Medicine. Any opinions, findings, and conclusions or recommendations expressed in this material are those of the author(s) and do not necessarily reflect the views of the National Science Foundation.

## Notes

### Competing Interest Statement

The authors have declared no competing interest.

