## Supplemental Figures and Tables for "A combined frequency domain near infrared spectroscopy and diffuse correlation spectroscopy system for comprehensive metabolic monitoring of inspiratory muscles during loading"

### Supplement

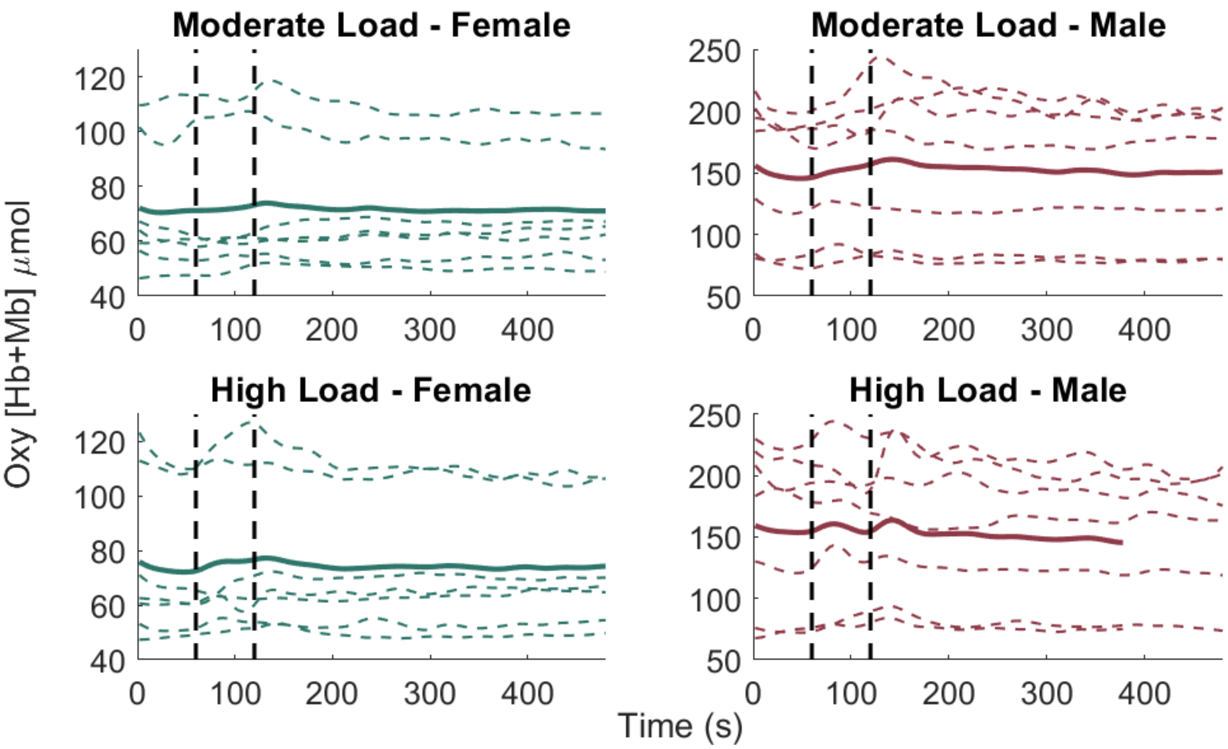


**Figure S1.** Filtered time traces of each subject and mean filtered time trace for each load and sex. Vertical black dashed lines indicate the start and end of the one-minute load portion of the respiratory exercise. The high load, male time trace has the last 100 seconds removed due to one subject coughing which resulted in large muscle activation.

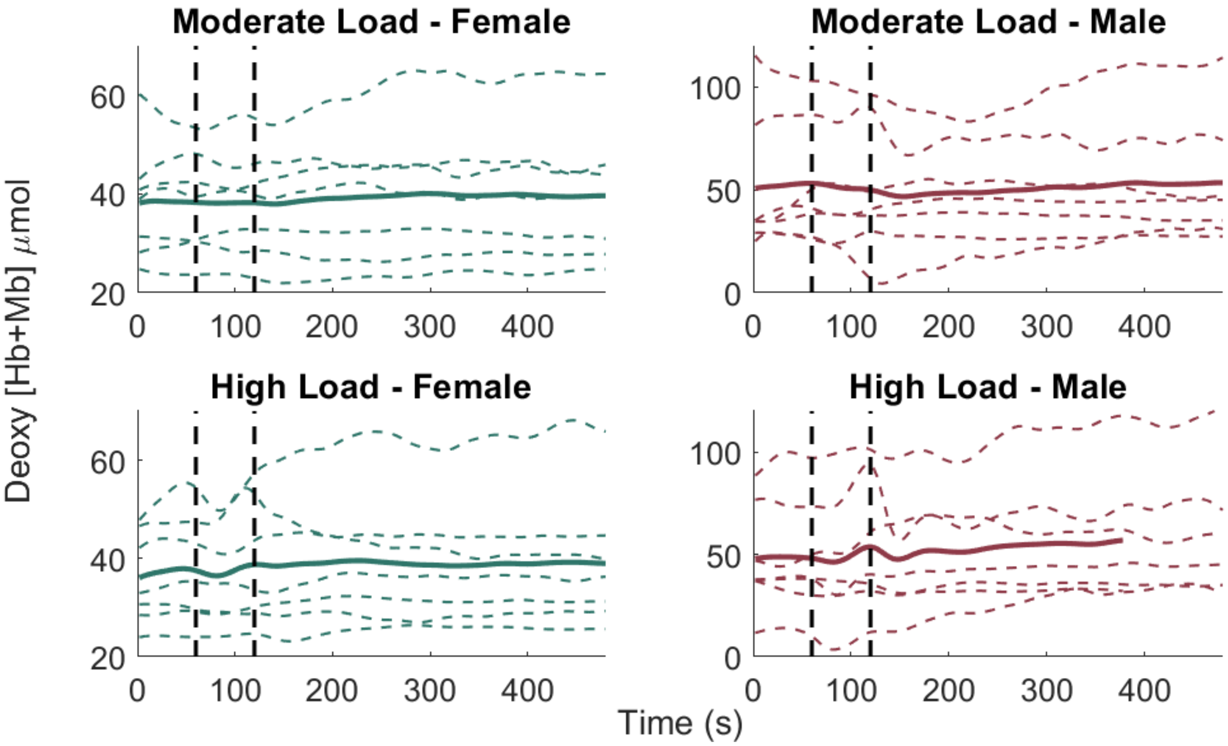


**Figure S2.** Filtered time traces of each subject and mean filtered time trace for each load and sex. Vertical black dashed lines indicate the start and end of the one-minute load portion of the respiratory exercise. The high load, male time trace has the last 100 seconds removed due to one subject coughing which resulted in large muscle activation.

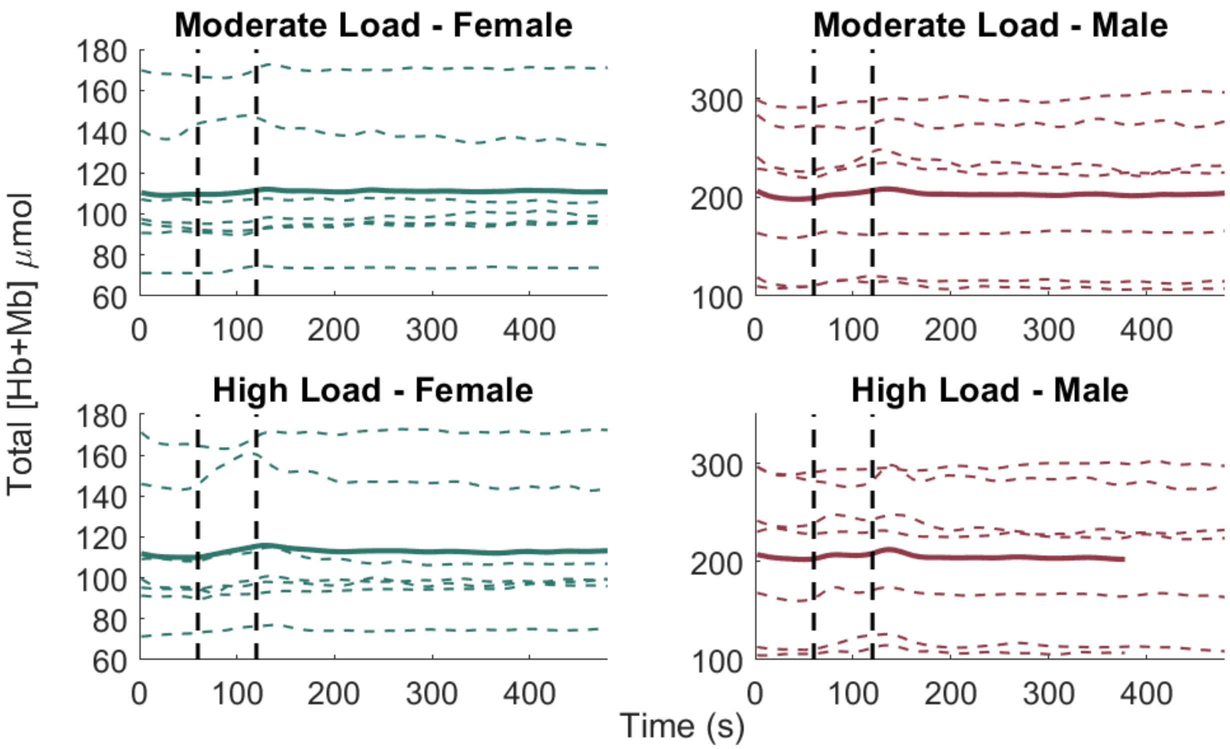


**Figure S3.** Filtered time traces of each subject and mean filtered time trace for each load and sex. Vertical black dashed lines indicate the start and end of the one-minute load portion of the respiratory exercise. The high load, male time trace has the last 100 seconds removed due to one subject coughing which resulted in large muscle activation.

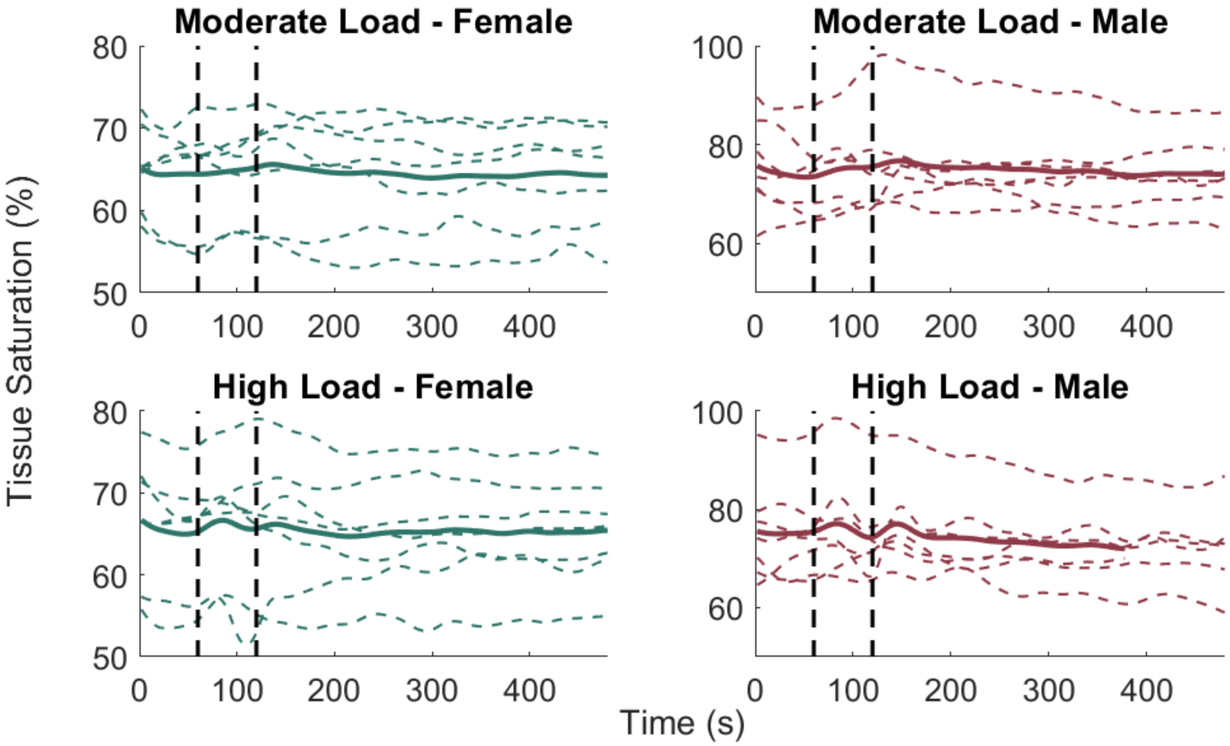


**Figure S4.** Filtered time traces of each subject and mean filtered time trace for each load and sex. Vertical black dashed lines indicate the start and end of the one-minute load portion of the respiratory exercise. The high load, male time trace has the last 100 seconds removed due to one subject coughing which resulted in large muscle activation.

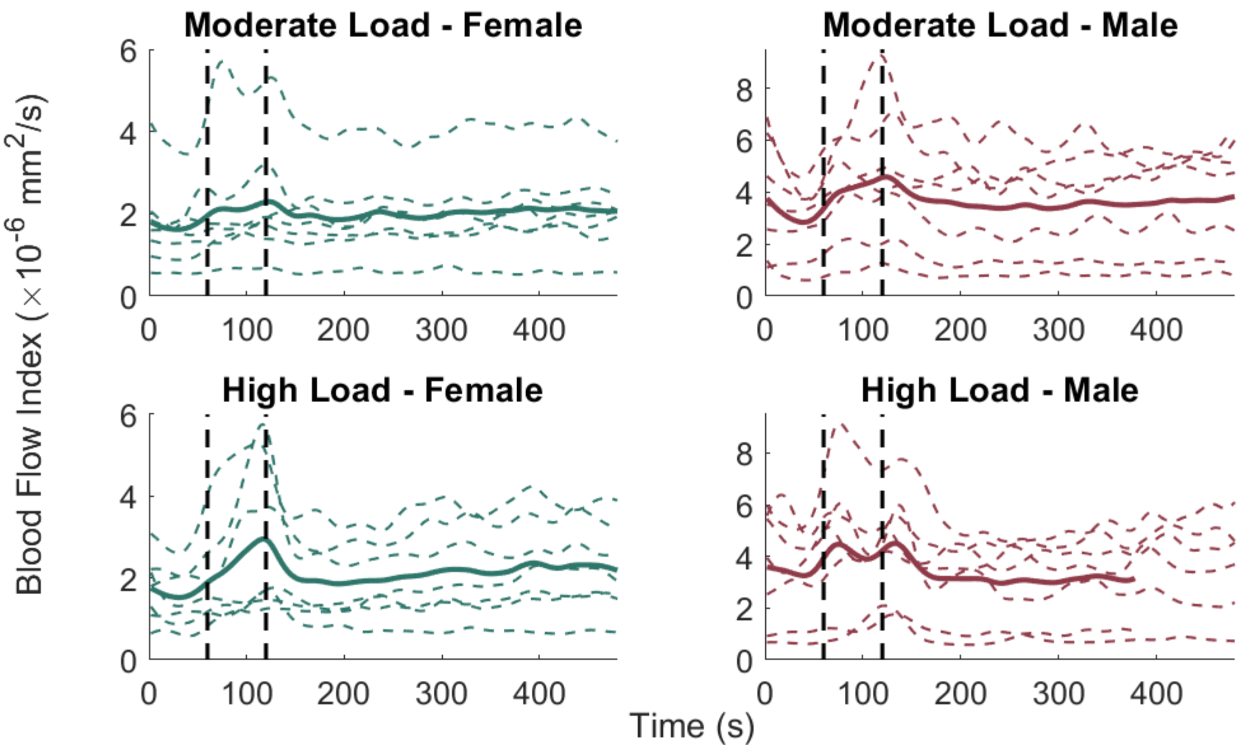


**Figure S5.** Filtered time traces of each subject and mean filtered time trace for each load and sex. Vertical black dashed lines indicate the start and end of the one-minute load portion of the respiratory exercise. The high load, male time trace has the last 100 seconds removed due to one subject coughing which resulted in large muscle activation.

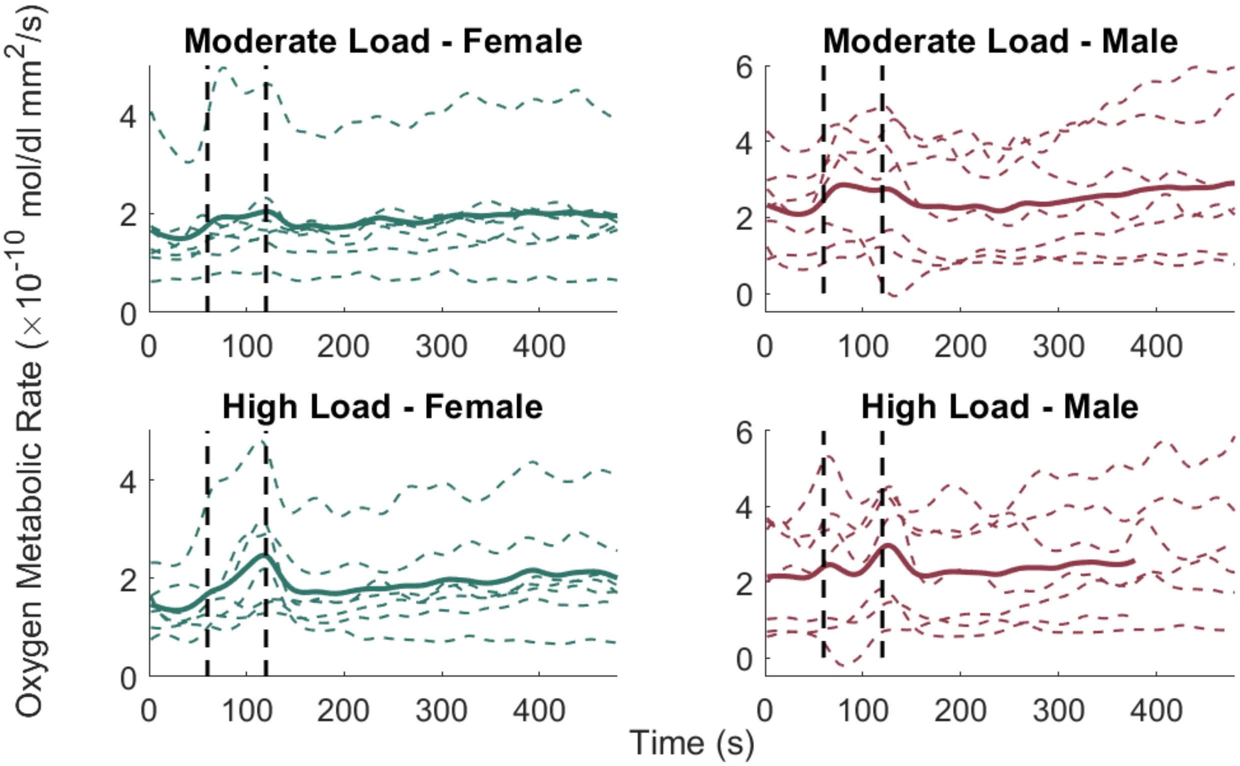


**Figure S1.** Filtered time traces of each subject and mean filtered time trace for each load and sex. Vertical black dashed lines indicate the start and end of the one-minute load portion of the respiratory exercise. The high load, male time trace has the last 100 seconds removed due to one subject coughing which resulted in large muscle activation.

**Table S1.** Mean + standard distribution of the offset (sec) of the six measured muscle parameters for all sex, load, and region of activation combinations. Values were calculated only if more than two peaks/valleys were present in the region of activation.

| | | Female | | Male | |
|---|---|---|---|---|---|
| | | 1st Region | 2nd Region | 1st Region | 2nd Region |
| Moderate Load | Oxy [Hb+Mb] | 17.3 ± 13.2 | 77.1 ± 40.6 | 23.3 ± 3.8 | 86.0 ± 31.8 |
| | Deoxy [Hb+Mb] | – | 58.3 ± 19.9 | 22.8 ± 8.1 | 118.0 ± 36.8 |
| | Total [Hb+Mb] | 27.3 ± 2.5 | 71.0 ± 8.0 | 23.6 ± 4.3 | 75.1 ± 4.5 |
| | $S_tO_2$ | 16.7 ± 15.2 | 73.7 ± 23.9 | 24.0 ± 1.6 | 87.3 ± 33.6 |
| | $BF_i$ | 16.0 ± 13.3 | 68.0 ± 18.1 | 19.7 ± 5.7 | 64.3 ± 5.8 |
| | $MRO_2$ | 14.8 ± 12.7 | 67.0 ± 17.8 | 17.4 ± 7.1 | 66.7 ± 7.2 |
| High Load | Oxy [Hb+Mb] | 25.3 ± 3.4 | 70.6 ± 10.5 | 22.0 ± 4.2 | 80.3 ± 3.7 |
| | Deoxy [Hb+Mb] | – | 89.6 ± 29.4 | 21.3 ± 4.1 | 107.6 ± 53.7 |
| | Total [Hb+Mb] | – | 72.9 ± 15.2 | – | 74.6 ± 8.1 |
| | $S_tO_2$ | 24.0 ± 3.3 | 84.4 ± 32.1 | 19.0 ± 4.6 | 105.7 ± 52 |
| | $BF_i$ | 20.0 ± 12.8 | 60.6 ± 5.5 | 12.3 ± 7.4 | 69.4 ± 6.7 |
| | $MRO_2$ | 22.7 ± 12.7 | 78.9 ± 47.2 | 12.7 ± 9.6 | 68.0 ± 6.0 |

**Table S2.** Mean + standard distribution of baseline and absolute change of the six measured muscle parameters for all sex, load, and region of activation combinations. Values were calculated only if more than two peaks/valleys were present in the region of activation.

| | | Female | | | Male | | |
|---|---|---|---|---|---|---|---|
| | | Baseline | 1st Region | 2nd Region | Baseline | 1st Region | 2nd Region |
| Moderate Load | Oxy [Hb+Mb] (µmol) | 71 ± 22 | 3.32 ± 2.96 | 3.76 ± 5.07 | 139 ± 50 | 6.13 ± 4.97 | 18.3 ± 15.7 |
| | Deoxy [Hb+Mb] (µmol) | 38 ± 10 | – | -0.90 ± 2.17 | 50 ± 29 | 1.75 ± 6.8 | -15.1 ± 13.0 |
| | Total [Hb+Mb] (µmol) | 109 ± 30 | -1.11 ± 0.85 | 3.74 ± 2.82 | 189 ± 69 | 3.72 ± 2.36 | 7.64 ± 3.23 |
| | $S_tO_2$ (%) | 64 ± 5.2 | 1.53 ± 0.38 | 1.49 ± 3.12 | 75 ± 7.9 | 2.64 ± 2.06 | 4.22 ± 5.12 |
| | $BF_i$ ($x10^{-6}$ $mm^2/s$) | 1.68 ± 0.93 | 0.72 ± 0.56 | 0.73 ± 0.56 | 2.82 ± 1.56 | 0.99 ± 0.44 | 1.81 ± 1.17 |
| | $MRO_2$ ($x10^{-10}$ mol/dl $mm^2/s$) | 1.55 ± 0.80 | 0.62 ± 0.53 | 0.54 ± 0.41 | 1.98 ± 1.02 | 0.73 ± 0.64 | 1.05 ± 0.66 |
| High Load | Oxy [Hb+Mb] (µmol) | 73 ± 25 | 1.76 ± 1.98 | 4.80 ± 7.04 | 144 ± 60 | 8.46 ± 10.2 | 13.7 ± 6.15 |
| | Deoxy [Hb+Mb] (µmol) | 37 ± 9.7 | – | 1.21 ± 3.56 | 47 ± 26 | -7.33 ± 2.48 | 8.85 ± 11.5 |
| | Total [Hb+Mb] (µmol) | 110 ± 31 | – | 5.94 ± 5.41 | 192 ± 72 | – | 7.87 ± 4.48 |
| | $S_tO_2$ (%) | 65 ± 7.1 | 1.82 ± 1.22 | 0.32 ± 4.33 | 74 ± 9.3 | 3.03 ± 3.41 | -1.33 ± 7.05 |
| | $BF_i$ ($x10^{-6}$ $mm^2/s$) | 1.60 ± 0.70 | 0.88 ± 0.73 | 1.40 ± 1.20 | 3.06 ± 1.84 | 1.33 ± 1.25 | 1.23 ± 1.12 |
| | $MRO_2$ ($x10^{-10}$ mol/dl $mm^2/s$) | 1.40 ± 0.50 | 0.99 ± 0.49 | 1.13 ± 0.73 | 1.96 ± 1.26 | 1.00 ± 0.40 | 0.86 ± 0.65 |

**Table S3.** Mean ± standard distribution of respiration rate (Hz) for all load and sex combinations during baseline and both regions of activation.

| | | Baseline | 1st Region | 2nd Region |
|---|---|---|---|---|
| Moderate Load | Female | 12.3 ± 1.2 | 11.7 ± 2.1 | 11.0 ± 0.8 |
| | Male | 13.1 ± 4.9 | 10.7 ± 2.1 | 10.4 ± 2.9 |
| High Load | Female | 10.9 ± 1.5 | 10.0 ± 2.1 | 10.7 ± 0.9 |
| | Male | 10.1 ± 3.3 | 9.3 ± 2.1 | 9.4 ± 2.0 |
